# Development of a 5-FU Modified miR-129 Mimic as a Therapeutic for Non-Small Cell Lung Cancer

**DOI:** 10.1101/2022.09.16.508317

**Authors:** Ga-Ram Hwang, John G. Yuen, Andrew Fesler, Hannah Farley, John D. Haley, Jingfang Ju

## Abstract

Lung cancer is the leading cause of cancer-related deaths in the United States with non-small cell lung cancer (NSCLC) accounting for most cases. Despite advances in cancer therapeutics, the 5-year survival rate has remained poor due to several contributing factors, including its resistance to therapeutics. Therefore, there is a pressing need to develop therapeutics that can overcome resistance. Non-coding RNAs, including microRNAs (miRNAs), have been found to contribute to cancer resistance and therapeutics by modulating the expression of several targets involving multiple key mechanisms. In this study, we investigated the therapeutic potential of miR-129 modified with 5-fluorouracil (5-FU) in NSCLC. Our results show that 5-FU modified miR-129 (5-FU-miR-129) inhibits proliferation, induces apoptosis, and retains function as a miRNA in NSCLC cell lines A549 and Calu-1. Notably, we observed that 5-FU-miR-129 was able to overcome resistance to tyrosine kinase inhibitors and chemotherapy in cell lines resistant to erlotinib or 5-FU. Furthermore, we observed that the inhibitory effect of 5-FU-miR-129 can also be achieved in NSCLC cells under vehicle-free conditions. Finally, 5-FU-miR-129 inhibited NSCLC tumor growth and extended survival *in vivo* without toxic side effects. Altogether, our results demonstrate the potential of 5-FU-miR-129 as a highly potent cancer therapeutic in NSCLC.

## Introduction

Lung cancer is the leading cause of cancer-related deaths in the United States. Lung cancer is generally classified into two main types, non-small cell lung cancer (NSCLC) and small cell lung cancer, with NSCLC accounting for ~85% of diagnosed cases.^1,2^ Several treatment strategies have been developed to treat patients with NSCLC, including chemotherapy, radiotherapy, and targeted therapy.^2^ For example, erlotinib is a tyrosine kinase inhibitor (TKI) used to treat patients diagnosed with NSCLC.^3–5^ TKIs, including erlotinib, gefitinib, and osimertinib, specifically inhibit activating epidermal growth factor receptors (EGFRs) mutations.^4–8^ Despite advancements in therapeutics to treat patients diagnosed with NSCLC, the 5-year survival rate after metastasis remains low (26%).^2,9^ One of the contributing factors for this is NSCLC’s resistance to cancer therapeutics.^3,10^ For example, certain mutations of EGFR, including T790M, have been found to contribute to resistance to first generation TKIs erlotinib and gefitinib.^11–13^ In addition to targeted therapeutics, resistance to chemotherapy in patients with NSCLC has also been well-documented.^3,10,14–17^ Therefore, there is a need to develop new therapeutics that can overcome resistance to cancer therapies in NSCLC.

The dysregulation of microRNA (miRNA) expression has been found to contribute to the pathogenesis of cancer and its resistance to therapeutics.^18–21^ MiRNAs are a class of small RNAs that are 21-25 nucleotides long and double-stranded. MiRNAs do not require complete complementary binding of their sequence with their mRNA targets, and instead, mature miRNAs bind to the 3’ untranslated region (UTR) of their targets via complementary binding with their seed sequence. Therefore, miRNAs can target and affect the expression of multiple different mRNA sequences.^22–24^ Because of their ability to downregulate multiple different targets, it has been reported that dysregulated expression of miRNAs can contribute to carcinogenesis, drug resistance, and stemness in several different cancers, including NSCLC.^23,25–28^ Specifically, expression of hsa-miR-129-5p (miR-129) has been reported to be downregulated in patients with NSCLC.^29–32^ Similarly, restoring miR-129 expression has been found to increase sensitivity to chemotherapy in NSCLC cells.^27^ These effects are, in part, due to the targets of miR-129, which include apoptosis regulator Bcl-2 (Bcl-2) and high mobility group protein B1 (HMGB1).^30,33–35^

In this study, we demonstrate the therapeutic potential of miR-129 modified with the chemotherapy drug 5-fluorouracil (5-FU) in NSCLC. By substituting uracil residues (U) within the guide strand sequence of miR-129, we have a developed a 5-FU modified miRNA mimic, 5-FU-miR-129 with the ability to be delivered to cells without the aid of a transfection vehicle. We observed that 5-FU-miR-129 inhibited cell proliferation, induced apoptosis, and was able to be delivered without the use of a transfection vehicle in NSCLC cell lines A549 and Calu-1. In addition, 5-FU-miR-129 retained miR-129’s target specificity and miRNA function. We also observed 5-FU-miR-129’s ability to overcome resistance to the EGFR TKI, erlotinib, as well as to 5-FU itself. We also identified that 5-FU-miR-129 has a pleiotropic effect on cell death genes, including *FAS* and *DPYSL4*, and death signaling pathways such as the NF-κB signaling pathway. Finally, we examined 5-FU-miR-129’s therapeutic potential compared to nucleoside analog chemotherapy drugs 5-FU and gemcitabine (Gem) *in vivo* and observed a significant inhibition of NSCLC tumor growth and an increase in median survival. As a result, we demonstrate that 5-FU-miR-129 has potential as a therapeutic agent that can overcome resistance to cancer therapeutics in NSCLC.

## Results

### 5-FU-miR-129 Inhibits Cell Proliferation, Induces Apoptosis and Cell Cycle Arrest in NSCLC

We investigated the efficacy of 5-FU-miR-129 on NSCLC cell lines A549 and Calu-1. As shown previously, we have developed 5-FU-miR-129 by substituting the uracil residues of miR-129’s guide strand with 5-FU (Figure 1A).^34^ We assessed the effects of 5-FU-miR-129 on cell proliferation in A549 and Calu-1 cells (Figure 1B & Figure S1). At 50 nM, 5-FU-miR-129 inhibited proliferation by 82.69% in A549 cells and by 68.31% in Calu-1 cells six days post-transfection (Figure 1B). In addition, the unmodified miR-129 was found to only inhibit proliferation in A549 cells (53.20%), but not in Calu-1 cells.

**Figure 1:**
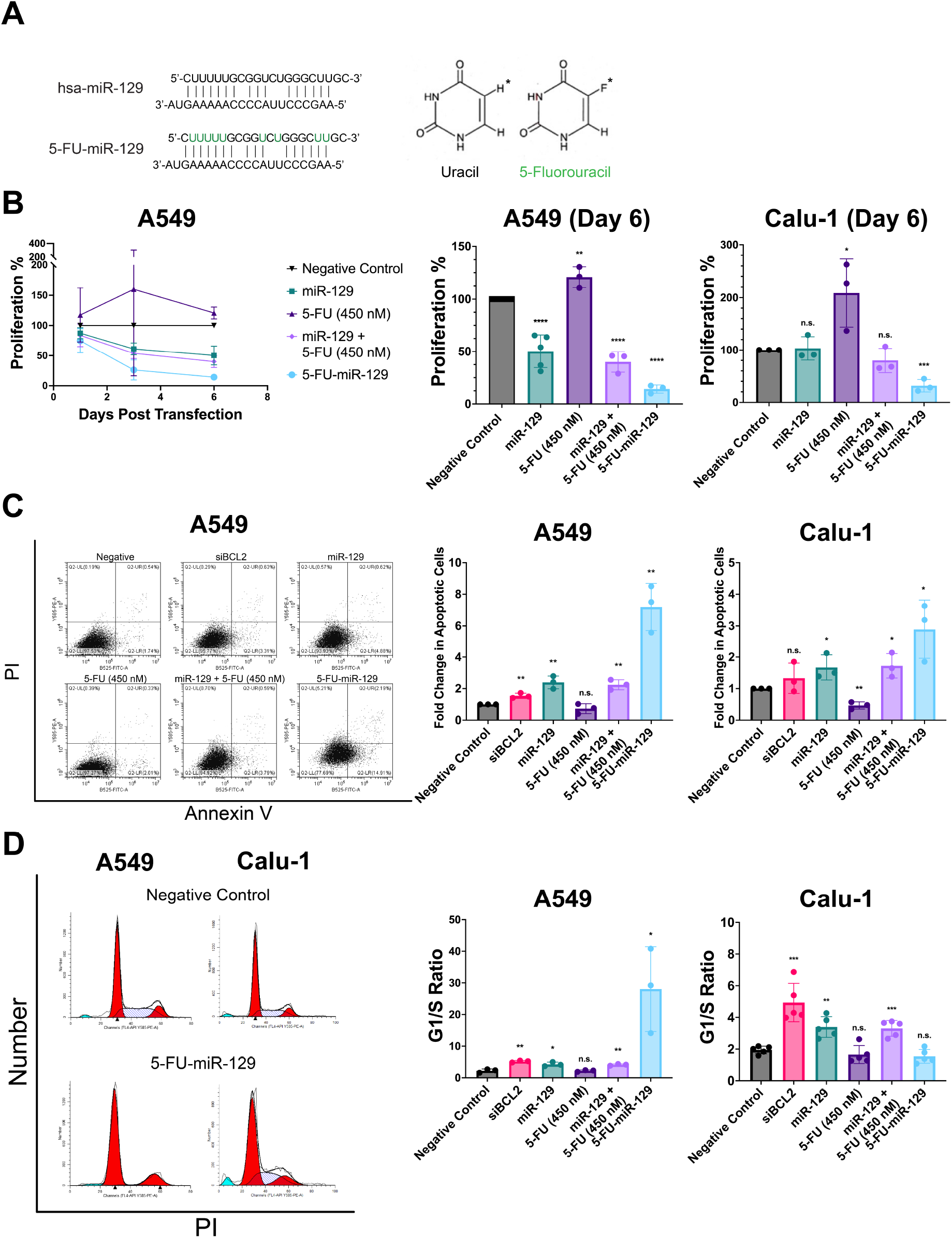
5-FU-Modified miR-129 Mimic Demonstrates Therapeutic Potential by Inhibiting Proliferation, Inducing Apoptosis and Inducing Cell Cycle Arrest in NSCLC *In Vitro*. (A) A 5-FU-modified miR-129 mimic (5-FU-miR-129) was developed by substituting the uracil residues within miR-129-5p’s sequence with 5-fluorouracil (green, 5-FU). 5-FU is a nucleoside analog of uracil with fluorine replacing the hydrogen on the fifth carbon of the uracil ring (*). (B) 5-FU-miR-129 inhibits proliferation in NSCLC *in vitro* 1, 3, and 6 days post-transfection in A549 cells. 1, 3, and 6 days post-transfection in Calu-1 cells is available in Figure S1. 6 days post-transfection, 5-FU-miR-129 inhibited cell proliferation by 82.69% in NSCLC cell lines A549 (*p* < 0.0001) and by 68.31% in Calu-1 (*p* = 0.0006). \**p*< 0.05, \*\**p*< 0.01, \*\*\**p*< 0.001, \*\*\**p*< 0.0001 (n = 3). (C) 5-FU-miR-129 induced apoptosis *in vitro* with a 7.2-fold increase in A549 cells *(p* = 0.0020) and a 2.9-fold increase in Calu-1 cells (*p* = 0.0240). \**p*< 0.05, \*\**p*< 0.01 (n = 3) (D) 5-FU-miR-129 induced cell cycle arrest in a cell-line dependent manner *in vitro*. Cell cycle arrest was only observed in A549 cells, as seen by a loss of cells in S phase and an increase in G1/S phase ratio in cells treated with 5-FU-miR-129 (28.05, *p* = 0.0288) compared to the negative control (2.211). A significant change in cell cycle arrest was not observed in Calu-1. \**p*< 0.05, \*\**p*< 0.01, \*\*\**p*< 0.001 (n=3). All transfections were performed with 50 nM of negative control, siRNA controls, unmodified miR-129, or 5-FU-miR-129. Cells were also treated with 450 nM of 5-FU to mimic the equivalent concentration of 5-FU in 5-FU-miR-129. Data are represented as mean ± SD.

We also assessed the effects of 5-FU-miR-129 on apoptosis and the cell cycle. 5-FU-miR-129 induced apoptosis in both NSCLC cell lines A549 and Calu-1 with a 7.2-fold and 2.9-fold increase of apoptotic cells, respectively (Figure 1C). In addition, unmodified miR-129 induced apoptosis with a 2.4 and 1.7-fold increase of apoptotic cells in A549 and Calu-1, respectively. Surprisingly, siBCL2 only induced an increase in apoptotic cells in A549 cells (1.5-fold increase). Furthermore, 5-FU-miR-129 induced an increase in the G1/S phase ratio in A549 cells, suggesting that 5-FU-miR-129 induces G1 cell cycle arrest in A549 cells (Figure 1D). In contrast, 5-FU-miR-129 did not induce a significant increase in the G1/S phase ratio in Calu-1 cells.

### 5-FU-miR-129 Downregulates Expression of Bcl-2 and HMGB1 in NSCLC

To demonstrate that 5-FU-miR-129 retains its function as a miRNA mimic of miR-129, we investigated 5-FU-miR-129’s ability to downregulate miR-129 targets Bcl-2 and HMGB1. *BCL2* and *HMGB1* have both been reported to be targets of miR-129.^30,33^ Therefore, we reasoned that 5-FU-miR-129 would also target *BCL2* and *HMGB1* (Figure 2A & 2C). In addition, 5-FU-miR-129 has previously been reported to downregulate Bcl-2 in colorectal cancer (CRC).^34^ 5-FU-miR-129 only downregulated Bcl-2 expression in A549 cells (Figure 2B). Unmodified miR-129 also downregulated Bcl-2 expression in both A549 and Calu-1 cells while 5-FU alone did not induce a significant change in Bcl-2 expression. The combination of unmodified miR-129 and 5-FU only downregulated Bcl-2 expression in A549 cells. In contrast, 5-FU-miR-129 downregulated HMGB1 expression in both A549 and Calu-1 cells (Figure 2D). Furthermore, HMGB1 expression was downregulated by miR-129, and the combination of unmodified miR-129 and 5-FU treatment conditions in both A549 and Calu-1 cells. Like its effect on Bcl-2, 5-FU was not found to have a significant effect on HMGB1 expression.

**Figure 2:**
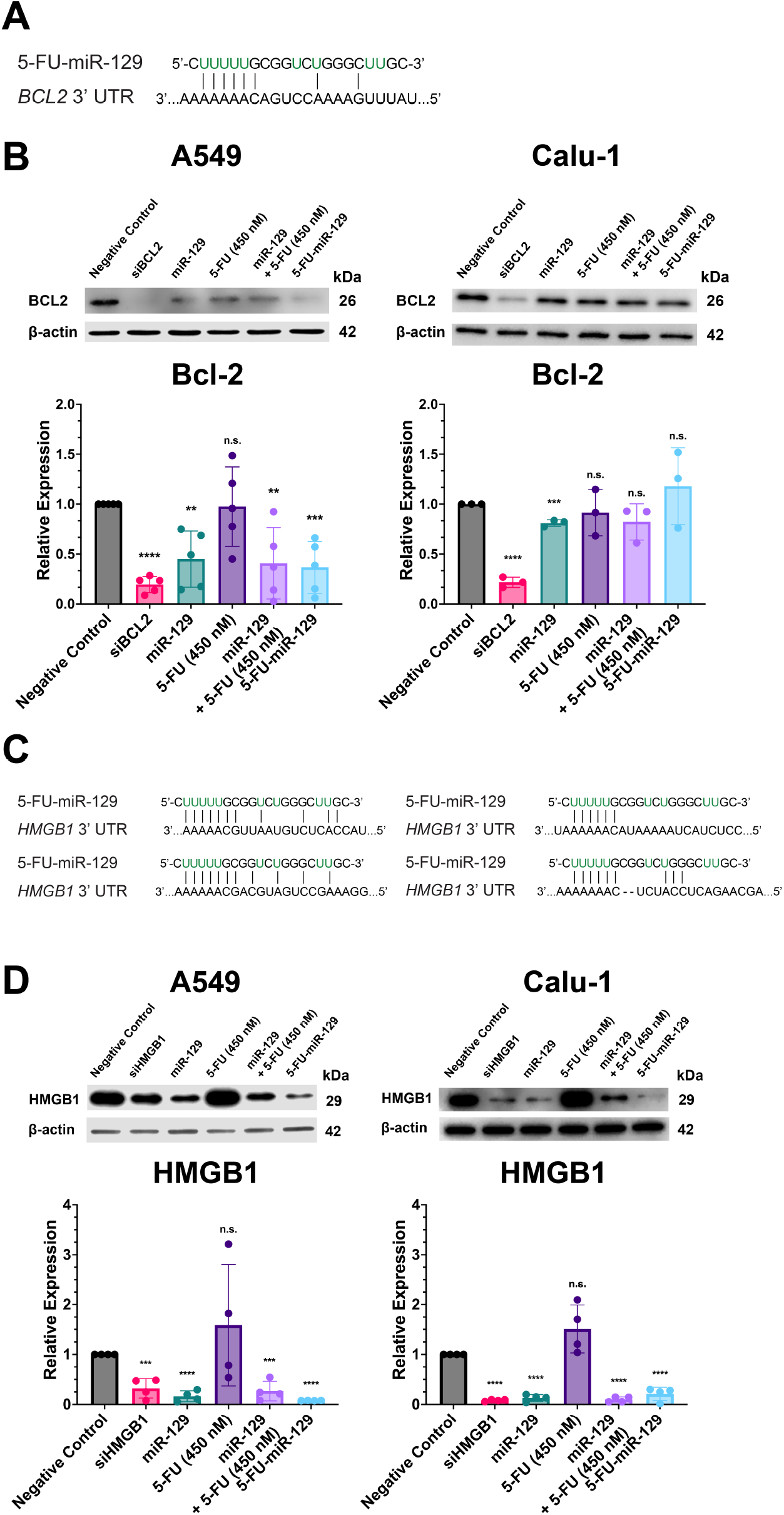
5-FU-miR-129 Retains miRNA Function and Knocks Down Expression of Bcl-2 and HMGB1. (A) 5-FU-miR-129 can potentially bind to miR-129’s predicted binding site within the 3’ UTR of *BCL2*. (B) 5-FU-miR-129 retains miRNA function as seen by a knock down of Bcl-2 expression in A549 cells (*p* = 0.0006). A knock down of Bcl-2 expression was not observed in Calu-1 cells. \*\**p*< 0.01, \*\*\**p*< 0.001, \*\*\*\**p*< 0.0001 (n = 3). (C) 5-FU-miR-129 can also potentially bind to miR-129’s four predicted binding sites within the 3’ UTR of *HMGB1*. (D) 5-FU-miR-129 was observed to knock down HMGB1 expression in both A549 (*p*< 0.0001) and Calu-1 cells (*p*< 0.0001). \*\*\**p*< 0.001, \*\*\*\**p*< 0.0001 (n = 4). All transfections were performed with 50 nM of negative control, siRNA controls, unmodified miR-129, or 5-FU-miR-129. Cells were also treated with 450 nM of 5-FU to mimic the equivalent concentration of 5-FU in 5-FU-miR-129. Data are represented as mean ± SD.

Due to the observed effects of 5-FU-miR-129 on cell cycle control only in A549 cells, we investigated the effects of 5-FU-miR-129 on the expression of cellular tumor antigen p53 (p53) and cyclin-dependent kinase inhibitor 1 (p21). 5-FU-miR-129 upregulated the expression of both p53 and p21 in A549 (Figure S2A). Treatment with siHMGB1 downregulated p53 expression while unmodified miR-129, or a combination of unmodified miR-129 and 5-FU, upregulated p21 expression. 5-FU alone did not have a significant effect on the expression of either p53 or p21 in both A549 and Calu-1 cells. Consistent with previous reports of a p53 deletion, ^36^ we did not observe expression of p53 (Figure S2B). Expression of p21 was also not observed in Calu-1 cells.

### 5-FU-miR-129 Promotes Autophagy in NSCLC in a Cell-Line Dependent Manner

Autophagy has been reported to have an intricate role in cell death, and autophagy has been reported be regulated by p21 and p53.^37–39^ Therefore, we investigated the effects of 5-FU-miR-129 on autophagy in NSCLC. A control with fresh serum (Negative Control (+Serum)) and a control under serum-starved conditions (Negative Control (Serum-Starved)) were used to represent inhibition and induction of autophagy, respectively (Figure 3A & 3B). Using the marker LC3B and the expression ratio between LC3B-II and LC3B-I to measure autophagy, 5-FU-miR-129 did not induce a significant change to autophagy in A549 cells (Figure 3A). In contrast, 5-FU-miR-129 promoted autophagy in Calu-1 cells (Figure 3B).

**Figure 3:**
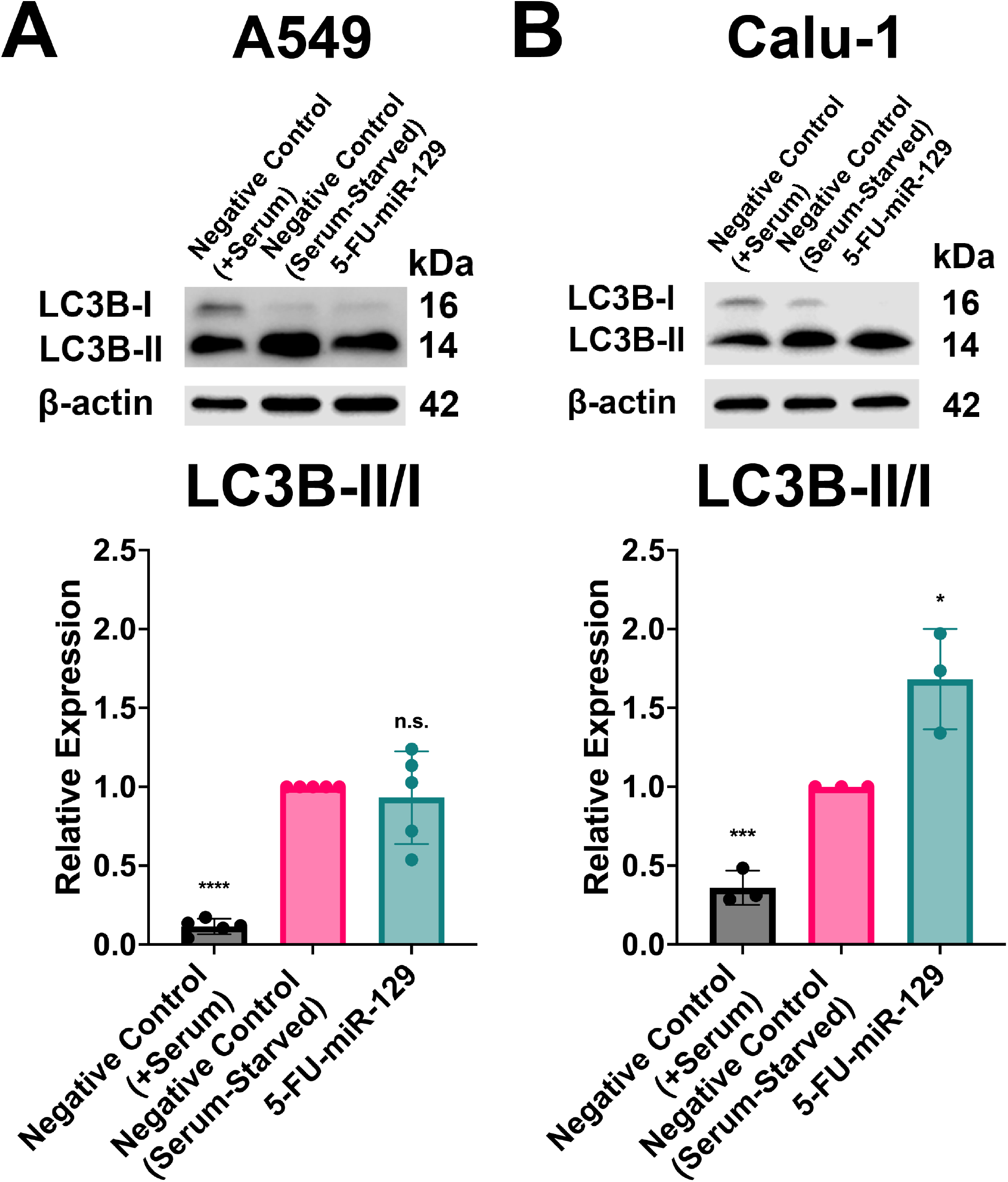
5-FU-miR-129 Affects Autophagy in a Cell-Line Dependent Manner in NSCLC *in vitro*. 4 hours prior to protein collection, cells were changed to serum-starved media (negative control and 5-FU-miR-129) to induce autophagy or changed to fresh media + serum (negative control + serum) to inhibit autophagy. (A) 5-FU-miR-129 did not have a significant effect on autophagy in A549 cells, as observed by a non-significant change to LC3B-II/I ratio. \*\*\*\**p*< 0.0001 (n = 5). (B) 5-FU-miR-129 was observed to promote autophagy in Calu-1 cells, as observed by a significant increase in LC3B-II/I ratio (*p* = 0.0255). \*\*\**p*< 0.001, \*\*\*\**p*< 0.0001 (n = 3). All transfections were performed with 50 nM of negative control and 5-FU-miR-129. Data are represented as mean ± SD.

### Vehicle-Free Delivery of 5-FU-miR-129 in NSCLC In Vitro

In this study, we investigated 5-FU-miR-129’s ability to be delivered without the aid of a transfection vehicle in NSCLC cells. Under vehicle-free conditions, 5-FU-miR-129 downregulated both Bcl-2 and HMGB1 in A549 cells (Figure 4A). The respective siRNAs (siBCL2 and siHMGB1), unmodified miR-129, and 5-FU did not induce a significant change in Bcl-2 or HMGB1 expression. Furthermore, we observed that vehicle-free transfection of 5-FU-miR-129 can inhibit cell proliferation. Six days post-transfection, 5-FU-miR-129 inhibited proliferation by 84.69% in A549 cells (Figure 4B).

**Figure 4:**
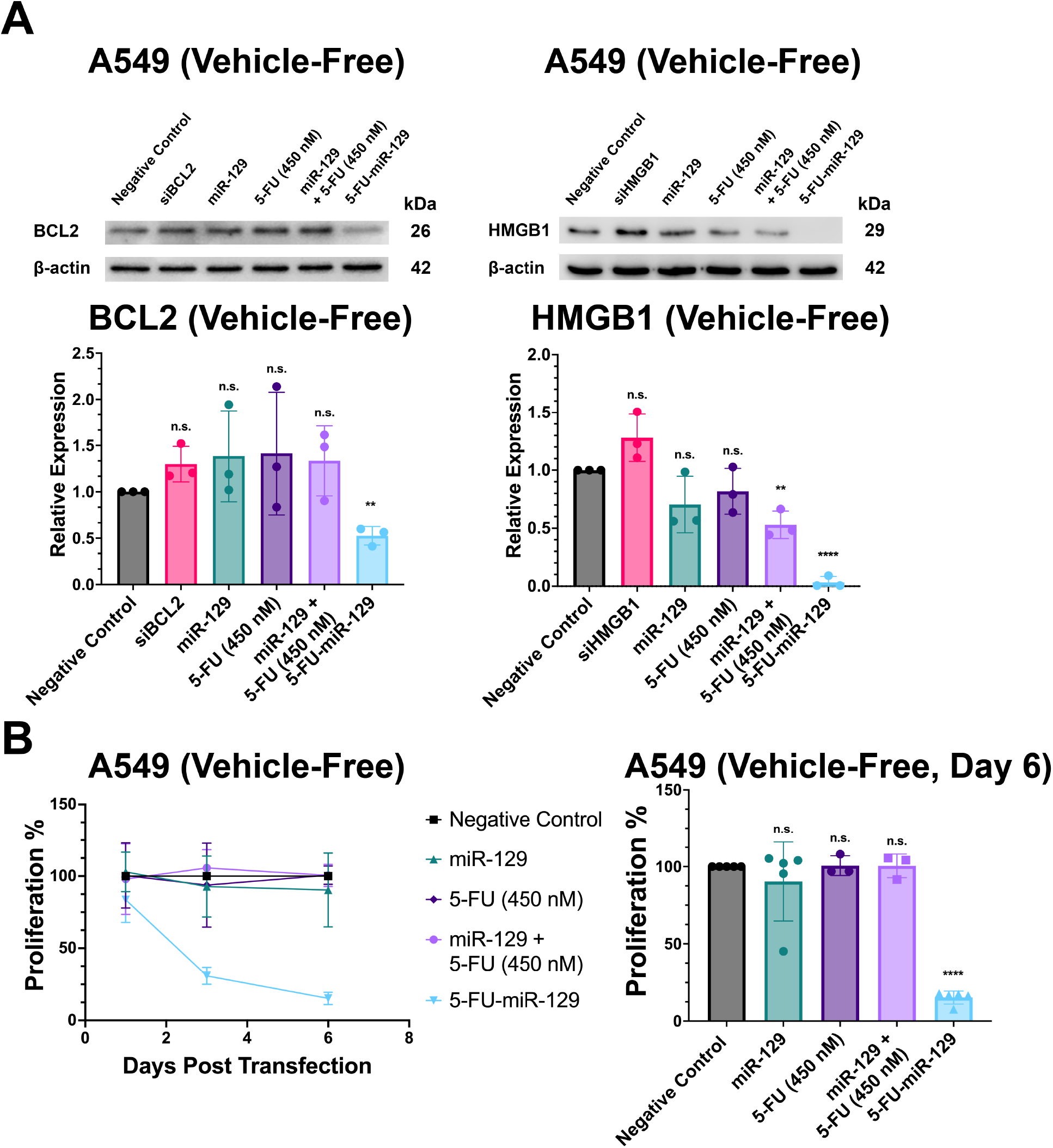
5-FU-miR-129 Can Be Transfected Vehicle-Free to NSCLC *In Vitro*. (A) 5-FU-miR-129 was found to be able knock down Bcl-2 (*p =* 0.0012) and HMGB1 (*p*< 0.0001) in A549 cells without the aid of a transfection vehicle. \*\**p*< 0.01, \*\*\*\**p*< 0.0001 (n = 3). (B) In addition, vehicle-free transfection of 5-FU-miR-129 was found to inhibit cell proliferation, as seen by an 84.69% reduction (*p*< 0.0001) in cell proliferation 6 days post-transfection via WST-1 assay. \*\*\*\**p*< 0.0001 (n = 3). All transfections were performed with 50 nM of negative control, siRNA controls, unmodified miR-129, or 5-FU-miR-129. Cells were also treated with 450 nM of 5-FU to mimic the equivalent concentration of 5-FU in 5-FU-miR-129. Data are represented as mean ± SD.

### 5-FU-miR-129 Overcomes Erlotinib and Chemoresistance in NSCLC

To demonstrate 5-FU-miR-129’s impact on resistance in NSCLC, we investigated the ability of 5-FU-miR-129 to inhibit proliferation in cells resistant to the EGFR TKI erlotinib. Using a cell line that is resistant to erlotinib, HCC827ER, we measured the IC_50_ of erlotinib in HCC827ER cells (64.26 nM), which is approximately 2.9 to 9.9-fold higher than previously reported values in its parental line (HCC827, 6.5 – 22 nM)^40,41^ and demonstrates HCC827ER’s resistance to erlotinib (Figure 5A). Compared to the IC_50_ of erlotinib (64.26 nM), we observed a 3.7-fold decrease in the IC_50_ of 5-FU-miR-129 (17.59 nM) in HCC827ER cells (Figure 5A). We then monitored 5-FU-miR-129’s ability to inhibit proliferation (Figure 5B). Six days post-transfection, 5-FU-miR-129 inhibited proliferation by 90.03% in HCC827ER cells at 50 nM. In addition, unmodified miR-129 inhibited proliferation by 27.67% in HCC827ER cells at 50 nM.

**Figure 5:**
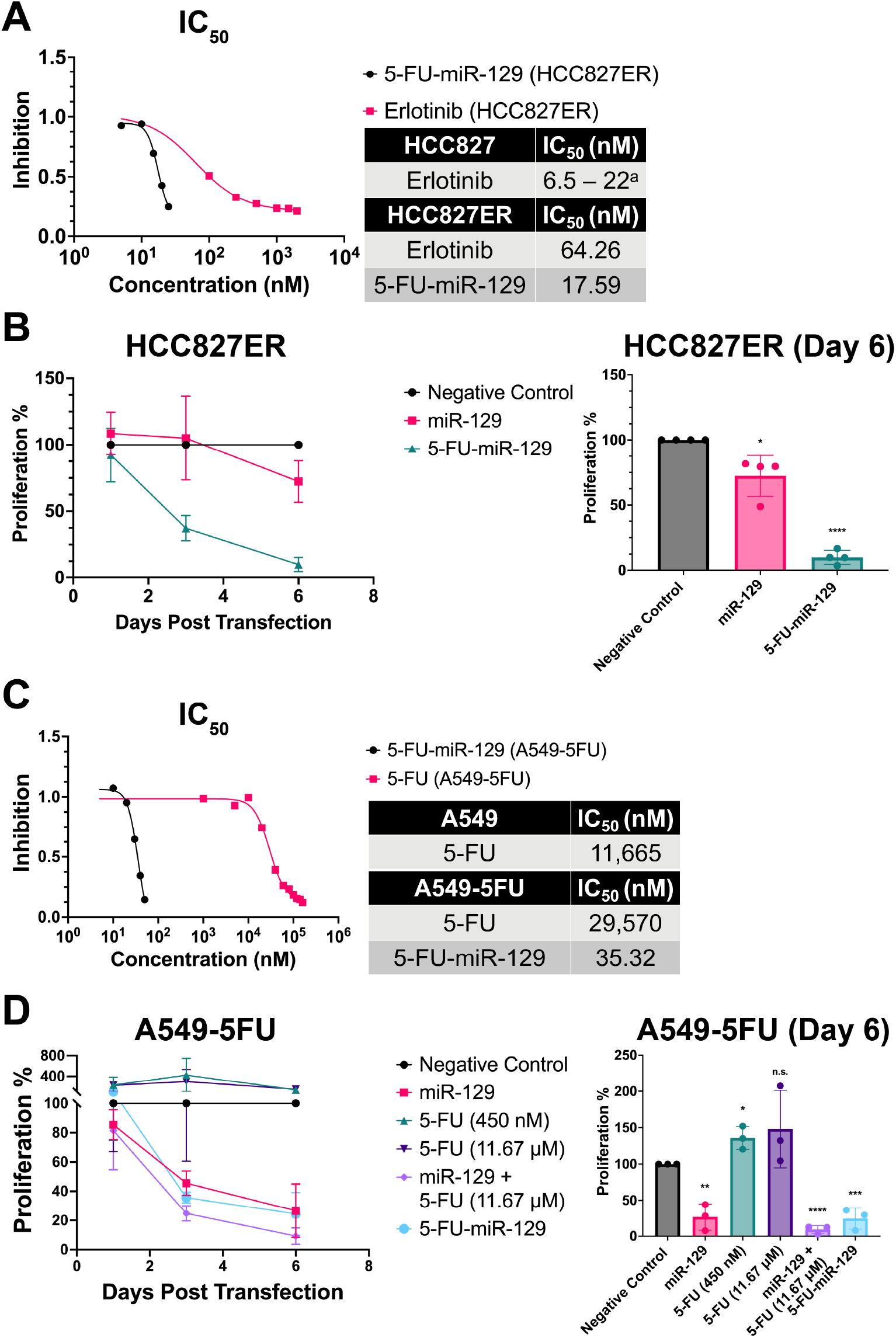
5-FU-miR-129 Can Overcome Resistance to Cancer Therapeutics in NSCLC *In Vitro*. (A) The IC_50_ values of erlotinib and 5-FU-miR-129 were measured in an NSCLC cell line that is resistant to erlotinib (HCC827ER). A 3.7-fold difference between the IC_50_ values of erlotinib and 5-FU-miR-129 was observed (n = 3). ^a^The IC_50_ of erlotinib in the parental line (HCC827) was determined by previously reported values.^40,41^ (B) 5-FU-miR-129 inhibits cell proliferation in HCC827ER. 6 days post-transfection, a 90.03% reduction in cell proliferation was observed in cells treated with 5-FU-miR-129 (*p*< 0.0001). All transfections were performed with 50 nM of negative control, miR-129, or 5-FU-miR-129. \**p*< 0.05, \*\*\*\**p*< 0.0001 (n = 4). (C) The IC_50_ values of 5-FU and 5-FU-miR-129 were measured in an NSCLC cell line that is resistant to the chemotherapy drug 5-FU (A549-5FU). An 837-fold difference between the IC_50_ values of 5-FU and 5-FU-miR-129 was observed (n = 3). (D) 5-FU-miR-129 inhibits cell proliferation in A549-5FU cells. 6 days post-transfection, a 75.56% reduction in cell proliferation was observed in cells treated with 5-FU-miR-129 (*p* = 0.0009). All transfections were performed with 50 nM of negative control, miR-129, or 5-FU-miR-129. A549-5FU cells were also treated with 5-FU at 450 nM or 11.67 μM to mimic the equivalent concentration of 5-FU in 5-FU-miR-129 or to mimic the observed IC_50_ of 5-FU in A549 to demonstrate resistance to 5-FU. \**p*< 0.05, \*\**p*< 0.01, \*\*\**p*< 0.001, \*\*\**p*< 0.0001 (n = 3). Data are represented as mean ± SD.

We also examined 5-FU-miR-129’s potency and ability to inhibit proliferation in cells resistant to 5-FU (A549-5FU). To confirm that A549-5FU acquired resistance to 5-FU, we measured the IC50 of 5-FU in both A549-5FU and A549 cells. The IC_50_ of 5-FU increased by 2.5-fold in A549-5FU cells (29.57 μM) compared to A549 cells (11.67 μM) (Figure 5C). We then measured the IC_50_ of 5-FU-miR-129 in A549-5FU cells and observed that the IC_50_ of 5-FU-miR-129 was 837-fold lower than the IC_50_ of 5-FU (29.57 μM). We then examined 5-FU-miR-129’s ability to inhibit proliferation in A549-5FU cells (Figure 5D). Six days post-transfection, 5-FU-miR-129 inhibited proliferation by 75.56% in A549-5FU cells at 50 nM. In addition, unmodified miR-129 inhibited proliferation by 73.37% and the combination of miR-129 and 5-FU inhibited proliferation by 90.77%. In contrast, at the IC_50_ of 5-FU in A549 cells (11.67 μM), 5-FU alone in did not induce a significant change in proliferation in A549-5FU cells.

All IC_50_ values were less than 50 nM in several NSCLC cells lines, demonstrating the efficacy of 5-FU-miR-129 at a low nM range (Table 1). We also compiled a table of IC_50_ values for the chemotherapy nucleoside analog agents 5-FU and Gem, and platinum-based agent cisplatin in the NSCLC cell line A549 and compared them to 5-FU-miR-129’s IC_50_ value (Table S1). The IC_50_ of 5-FU-miR-129 (13.33 nM) was 37.4-fold and 853-fold less than the IC_50_ of Gem (458.48 nM)^42^ and cisplatin (11.37 μM), respectively. Our results show that 5-FU-miR-129 is at least 30-fold more effective than chemotherapy agents.

**Table 1:**
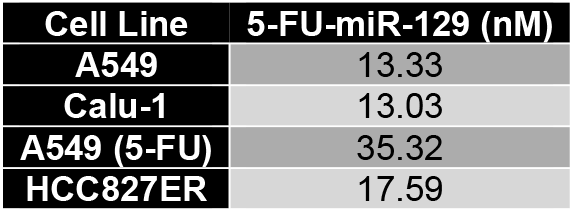
IC_50_ Values of 5-FU-miR-129 in NSCLC Cell Lines.

### Identifying Changes in mRNA Expression of Cell Death Pathway Induced by 5-FU-miR-129

We investigated changes in the global impact of cell death genes and pathways via RNA expression analysis after treatment with 5-FU-miR-129 in A549 (Table S2) and Calu-1 cells (Table S3). Of note, *ULK1* was found to be consistently downregulated in both A549 and Calu-1. Consistent with our western blot data, we also observed downregulated expression of *BCL2* only in A549, but not in Calu-1 cells. In contrast, *BCL2A1, CYLD, DPYSL4, FAS, and GADD45A* were found to be consistently upregulated in both A549 and Calu-1 cells. Using GeneSpring GX (Agilent), direct interactions between upregulated and downregulated targets were analyzed (Figure 6A & 6C). Pathway analysis of identified targets was also performed via Enrichr using the KEGG 2021 Human Database (Figure 6B & 6D).^43–45^ Of note, changes in both the apoptosis and NF-κB signaling pathway were found to be associated with 5-FU-miR-129 in A549 and Calu-1 cells.

**Figure 6:**
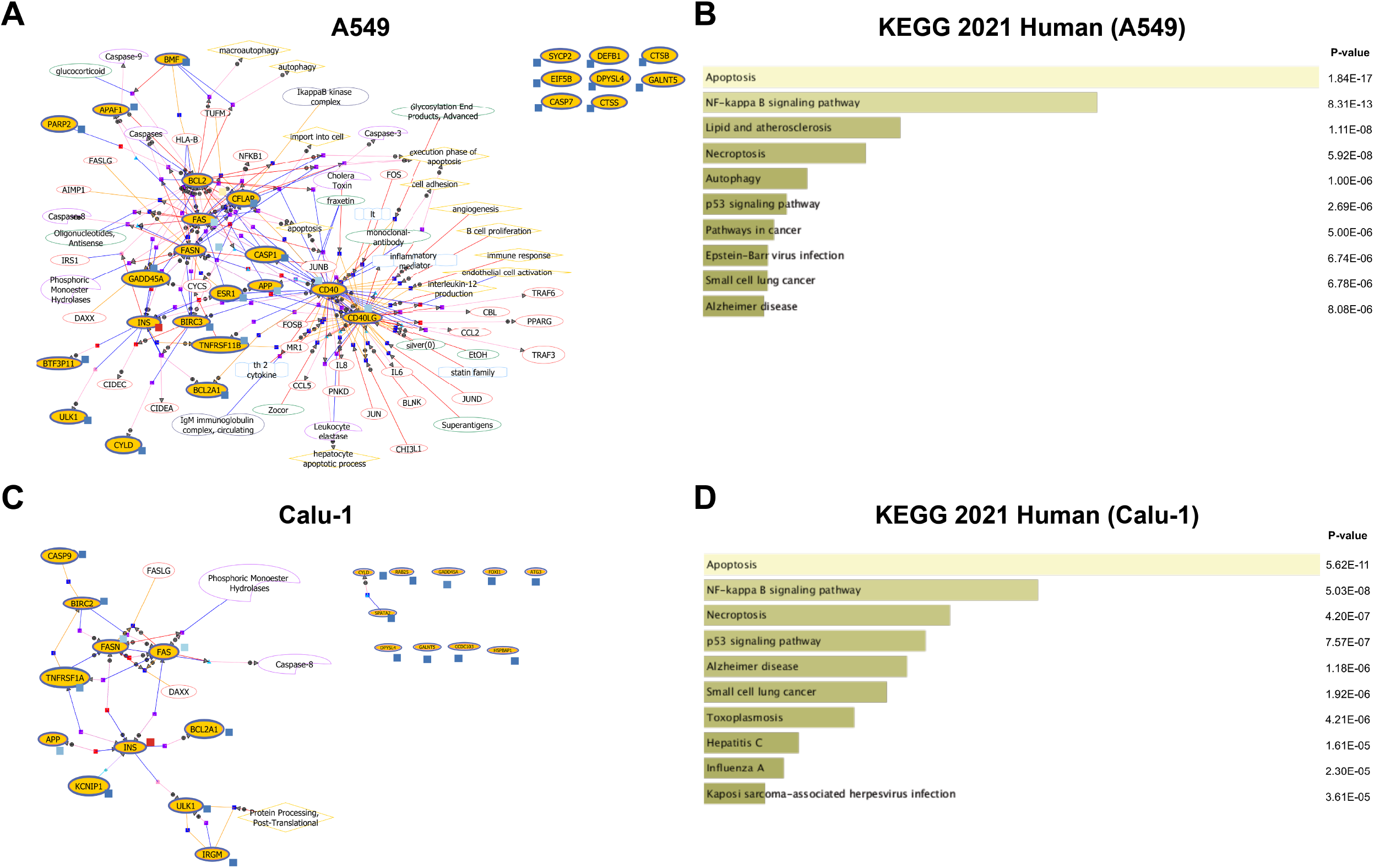
5-FU-miR-129 Affects Global mRNA Expression in Targets Related to Cell Death. NSCLC cell lines A549 and Calu-1 were transfected with either a negative control or 5-FU-miR-129. Extracted RNA per condition were pooled together (n = 3) and used in QIAGEN’s RT^2^ Profiler Array to identify changes in mRNA expression in targets related to cell death via RT-PCR. (A, B) Changes in mRNA expression were identified in A549 cells and the results were analyzed and visualized for pathway analysis via GeneSpring GX and Enrichr. (C, D) Changes in mRNA expression were also identified in Calu-1 cells and the results were analyzed and visualized for pathway analysis via GeneSpring GX and Enrichr.

### Therapeutic Potential of 5-FU-miR-129 in NSCLC In Vivo

To demonstrate the therapeutic potential of 5-FU-miR-129 in NSCLC *in vivo*, we examined the effects of 5-FU-miR-129 on a luciferase-expressing NSCLC cell line (A549-Luc) in NOD-SCID mice. Compared to the vehicle control (negative) group, decreased luciferase expression was observed in mice treated with 5-FU-miR-129 (Figure 7A). By day 37, a significant decrease in tumor growth was observed in mice treated with 5-FU-miR-129 (Figure 7B & 7C). Compared to the increase in tumor growth in the negative control group (13.22-fold growth), 5-FU-miR-129 induced a greater than 4.9-fold reduction in tumor growth in mice (2.67-fold growth, *p* = 0.0008).

**Figure 7:**
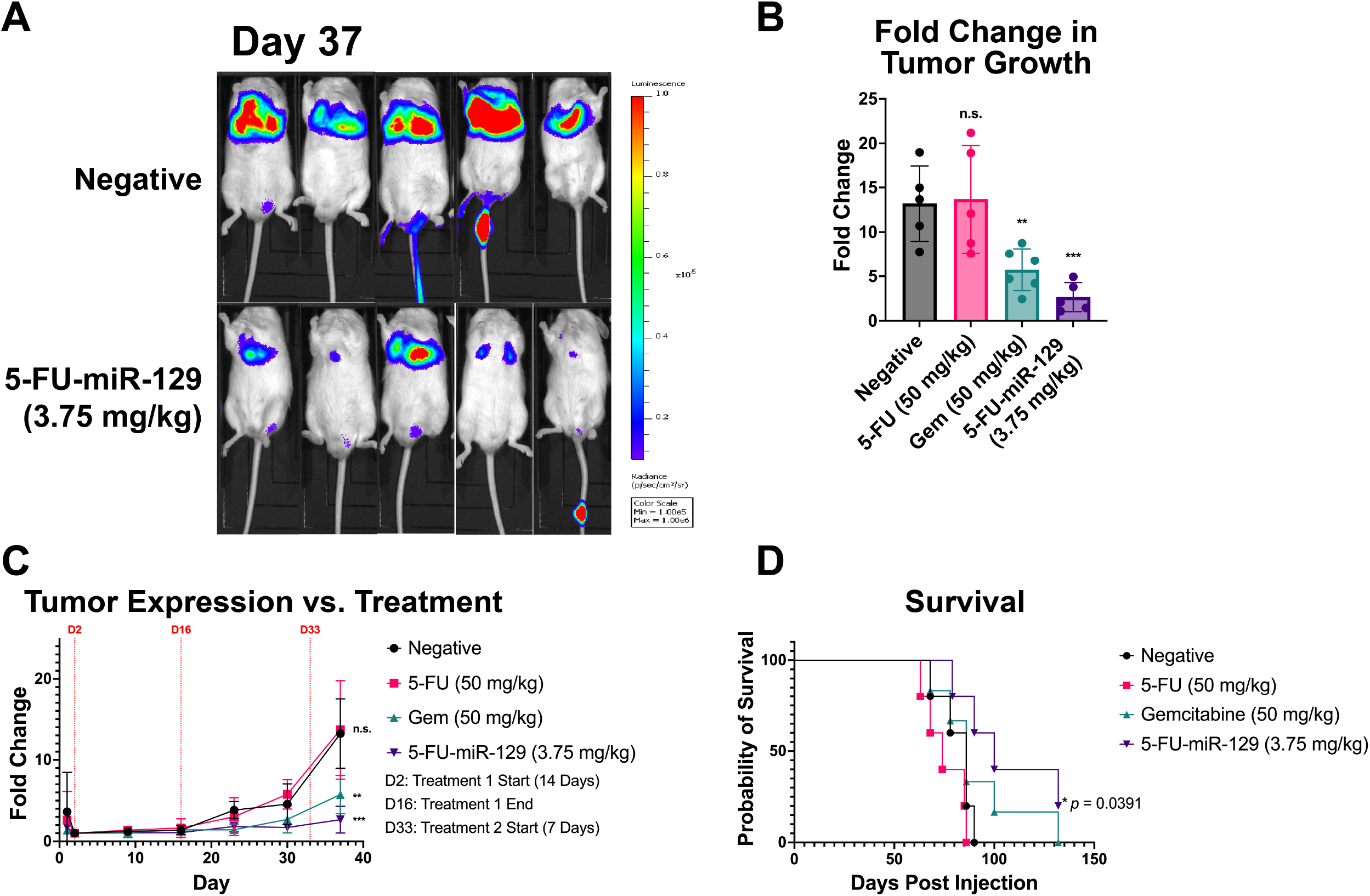
5-FU-miR-129 Can Inhibit NSCLC Tumor Growth *In Vivo* and Promote Survival. A luciferase-expressing line of NSCLC (A549-Luc) was injected (1 x 10^7^ cells per mouse) into NOD-SCID mice via intravenous tail vein injection. Post-tumor injection, mice were given two rounds of treatment (D2-16 and D33-40) with either vehicle control (negative), 5-FU (50 mg/kg), gemcitabine (Gem) (50 mg/kg), or 5-FU-miR-129 (3.75 mg/kg) via intravenous tail injection (negative and 5-FU-miR-129) or intraperitoneal injection (IP) (5-FU and Gem). (A) Tumor growth was monitored via luciferase expression using the IVIS system as seen by IVIS images of the negative and the 5-FU-miR-129 treated group. IVIS images of 5-FU and Gem treated groups are available in Figure S3. (B, C) 37 days post-injection of tumor cells, 5-FU-miR-129 (*p* = 0.0008) and Gem (*p* = 0.0049) inhibited NSCLC tumor growth when compared to the negative control group. \*\**p*< 0.01, \*\*\**p*< 0.001 (n = 5). (D) Mice were also monitored for survival, and only mice treated with 5-FU-miR-129 were found to have extended survival from 86 days to 100 days (HR = 0.3209, *p* = 0.0391). Data are represented as mean ± SD.

In addition, Gem decreased tumor growth (5.74-fold, *p* = 0.0049), but 5-FU did not induce a significant change in tumor growth in mice. We also monitored and compared survival between the different treatment groups (Figure 7D). Compared to the negative control group, only 5-FU-miR-129 induced an increase in the median survival for mice from 86 days to 100 days (*p* = 0.0391, hazard ratio = 0.3209). Hair loss and significant loss in body weight were also not observed in mice treated with 5-FU-miR-129 (data not shown).

## Discussion

In this study, we developed 5-FU-miR-129 as a potential effective cancer therapeutic in NSCLC. This is highly consistent with our recent findings that 5-FU-modified miRNAs can serve as a platform miRNA-based therapeutic technology in several different cancer types, including the therapeutic potential of 5-FU-miR-129 in CRC.^34,46–49^ As seen by its ability to inhibit proliferation and induce apoptosis in NSCLC cell lines A549 and Calu-1 (Figure 1), our results demonstrate 5-FU-miR-129’s potential in NSCLC.

Our results also suggest that 5-FU-miR-129 retains its function as a miRNA, specifically, its ability to pleiotropically knock down expression of miR-129 targets, HMGB1 and Bcl-2. Our previous studies have demonstrated that miR-129 and 5-FU-miR-129 can downregulate Bcl-2 in CRC ^33,34^. Furthermore, HMGB1 and Bcl-2 both have been reported to be promising targets in cancer therapeutics, and both have been reported to promote autophagy and resistance to chemotherapy in NSCLC.^30,50–55^ Bcl-2 is a well-studied protein that has been reported to inhibit apoptosis,^56^ and several Bcl-2 inhibitors have been developed.^57^ We also observed that 5-FU-miR-129 did not downregulate Bcl-2 expression in Calu-1 cells (Figure 2B), which may be due to a difference in the abundance of Bcl-2 mRNA transcript levels. Although Bcl-2 was not knocked down in Calu-1 cells, we observed that 5-FU-miR-129 induced a significant increase in apoptotic cells in both A549 and Calu-1. This is not surprising as miRNA targets are dependent on the cellular context.^58^ Furthermore, despite observing a knock down of Bcl-2 via siBCL2 (Figure 2B), siBCl2 did not induce an increase in apoptotic cells (Figure 1C). This suggests that 5-FU-miR-129 may induce apoptosis through a pathway that may not be dependent on Bcl-2. We also reasoned that knock down of HMGB1 by 5-FU-miR-129 would impact autophagy. Surprisingly, we observed that knock down of HMGB1 by 5-FU-miR-129 did not affect autophagy in A549 cells and increased autophagy in Calu-1 cells. 5-FU-miR-129’s dependence on cellular context for target selection may also influence 5-FU-miR-129’s effect on autophagy.

To better understand the impact of 5-FU-miR-129 in NSCLC cells, we quantified changes in mRNA expression of genes and pathways related to cell death mechanisms. These results are highly consistent with the pleiotropic effects of 5-FU-miR-129 in NSCLC in triggering apoptosis and autophagy. *FAS* and *GADD45A* have been reported to play a role in inducing apoptosis and have also been previously identified to have a positive correlation with patient prognosis in patients with NSCLC.^59–63^ *DPYSL4* has been reported to function as a tumor suppressor,^64^ and interestingly, *DPYSL4* has also been reported to play a role in inducing necrosis.^65^ However, necrosis was not an area of focus for this study, and 5-FU-miR-129’s role in necrosis can be examined in future studies. Consistent with our western blot results, *BCL2* was observed to only be downregulated in A549, but not in Calu-1 cells. Furthermore, pathway analysis of the observed changes in mRNA expression suggests that 5-FU-miR-129 primarily affects the apoptosis pathway in both A549 and Calu-1 cells (Figure 6B & 6D). In addition, the NF-κB signaling pathway was suggested to be affected by 5-FU-miR-129, which has been reported to play a major role in inflammation and cancer, and HMGB1 has also been reported to activate this pathway.^66–68^ Therefore, examining 5-FU-miR-129’s role in the NF-κB signaling pathway may be promising. 5-FU-miR-129’s effect on global changes in mRNA expression and protein abundance can also be further elucidated with RNA sequencing and proteomic studies.

Currently, the development and refinement of oligonucleotide delivery has been an area of study in RNA-based therapeutics.^46,69^ The inherent toxicity and unforeseen side effects of some delivery vehicles have been an obstacle in the effective delivery of RNA-based therapeutics.^46,70,71^ Our results demonstrate that 5-FU-miR-129 can be delivered to NSCLC cell without the aid of a delivery vehicle via knock down of Bcl-2 and HMGB1 expression and inhibition of proliferation (Figure 4). This is consistent with our previous studies that have shown that 5-FU-modified miRNAs can be delivered without the use of a transfection vehicle in pancreatic, gastric, lung, and breast cancer.^46–49^ In addition, this is consistent with previous studies that show that 5-FU-miR-129, specifically, inhibits proliferation in CRC via vehicle-free delivery.^34^ Notably, 5-FU is not listed among the list of current treatments for patients with NSCLC.^72^ Although 5-FU is not currently used, our results demonstrate that the 5-FU modification is necessary for miR-129 to enter cells without the aid of a delivery vehicle. Because the inhibitory effect of 5-FU-miR-129 can be achieved under vehicle-free conditions, we believe that this feature can overcome a key obstacle of RNA-based therapeutics.

This study also revealed cell line-dependent differences that were not observed in previous studies. Unlike previous studies, 5-FU-miR-129 was shown to have different effects on cell cycle arrest, autophagy, and knock down in miR-129 targets between A549 and Calu-1 cells. We hypothesize that several factors could have contributed to the differences observed in our study. A549 expresses wild-type p53 and Calu-1 is a p53-null mutant. This can lead to a difference of expression of downstream targets of p53, as demonstrated by a difference in expression of p21 between A549 and Calu-1 cells. Therefore, this difference can also affect pathways related to cell death and survival, including cell cycle arrest and autophagy.^34,38,73,74^ Interestingly, this could also contribute to the difference in induced apoptotic cells observed between A549 and Calu-1 cells due to the induction of autophagy in Calu-1 cells. Furthermore, although A549 and Calu-1 are both classified as NSCLC, A549 is also classified under the adenocarcinoma subtype while Calu-1 is classified under the squamous cell carcinoma subtype. Despite both subtypes being classified as NSCLC, the difference in subtype can contribute to differences in clinical decisions, including treatment.^75,76^ Therefore, 5-FU-miR-129’s relationship with p53 and NSCLC subtypes should be investigated in future studies. However, despite the different genetic backgrounds in NSCLC cells, we demonstrate that 5-FU-miR-129 inhibits proliferation, induces apoptosis, retains miRNA function, and can be delivered vehicle-free in NSCLC *in vitro*.

Importantly, as a potential therapeutic for NSCLC that can overcome resistance to cancer therapeutics, we demonstrate that 5-FU-miR-129 can overcome resistance to chemotherapy and TKIs. EGFR TKIs are currently being used in a clinical setting for patients diagnosed with NSCLC, but patients commonly develop resistance to TKIs.^3–5,11,75,76^ Our results demonstrate that 5-FU-miR-129 can provide a foundation in using 5-FU-miR-129 to overcome erlotinib resistance in the development of future therapeutics for NSCLC. In addition to TKIs, *KRAS* inhibitors targeting the G12C mutation were approved for treating NSCLC.^77,78^ A549 and Calu-1 are NSCLC cell lines with *KRAS* mutations, specifically G12S (A549) and G12C (Calu-1) with G12C being the most common *KRAS* mutant variant in NSCLC.^79^ However, resistance to these inhibitors have also been observed.^80,81^ *KRAS* is one of the predicted targets of miR-129 found through the target prediction software, TargetScan (data not shown), and we reason that 5-FU-miR-129 may also have the potential to overcome resistance to *KRAS* inhibitors.^82^ In addition to targeted inhibitors, platinum-based chemotherapies, including cisplatin, are used in a clinical setting for patients diagnosed with NSCLC.^83^ Compared to chemotherapy agents 5-FU, Gem, and cisplatin, 5-FU-miR-129 demonstrates potential as an efficacious therapeutic by exhibiting nanomolar potency and overcoming resistance to chemotherapy.

We also demonstrate 5-FU-miR-129’s promising potential as a cancer therapeutic in NSCLC within an *in vivo* context. Compared to the negative control group and 5-FU alone group, 5-FU-miR-129 was able to inhibit NSCLC tumor growth and increase survival (Figure 7). In addition, we did not observe any noticeable declines in animal health (e.g., hair loss and body weight loss; data not shown), thus suggesting 5-FU-miR-129’s potential to avoid side effects associated with cancer therapeutics. Interestingly, the mice in this study were treated with 5-FU-miR-129 at 3.75 mg/kg per injection. The 5-FU-miR-129 treated group was given a dose that is over 13-fold less than the 5-FU (50 mg/kg) and Gem (50 mg/kg) treated groups. This suggests that 5-FU-miR-129 is also effective at a lower dose than other chemotherapy drugs *in vivo*.

In summary, our studies demonstrate 5-FU-miR-129’s therapeutic potential in NSCLC by inhibiting cell proliferation, induces apoptosis, and can overcome resistance to both chemotherapy and targeted therapy. Our studies demonstrate that 5-FU-miR-129 exhibits high potency compared to therapies that are currently in clinical use for NSCLC. We demonstrate that 5-FU-miR-129 impacts promising targets Bcl-2 and HMGB1, along with other critical genes and pathways related to cell death. 5-FU-miR-129 is also able to be delivered to NSCLC cell without the aid of a delivery vehicle, a unique feature for nucleic acid-based medicine to reduce potential toxicity associated with the delivery vehicle. Finally, we also demonstrate the proof-of-concept that 5-FU-miR-129 can inhibit NSCLC tumor growth *in* vivo while increasing survival without toxic side effects. This study establishes the foundation to further develop 5-FU-miR-129 as an effective therapeutic for treating NSCLC.

## Materials and Methods

### Cell Lines and Culture

Human non-small cell lung cancer cell lines (A549 and Calu-1) were purchased from the American Type Culture Collection (ATCC, Manassas, VA, USA) and maintained in Ham’s F-12K (Kaighn’s) Medium or McCoy’s 5A (Modified) Medium (Thermo Fisher Scientific, Waltham, MA, USA) supplemented with 10% fetal bovine serum (FBS) (MilliporeSigma, St. Louis, MO, USA). FBS was sterilized via Steriflip-GP, 0.22 μm (MilliporeSigma, St. Louis, MO, USA) prior to addition to media.

The resistant cell line A549 (5-FU) was generated by treating the A549 cell line with 5-fluorouracil (MillporeSigma, St. Louis, MO, USA) starting at the IC_12.5_ for 48 hours in Ham’s F-12K (Kaighn’s) Medium supplemented with 10% dialyzed fetal bovine serum (DFBS) (Thermo Fisher Scientific, Waltham, MA, USA). After 48 hours, media was changed to fresh media until plates recovered from treatment. This process was repeated with increasing concentrations 5-fluorouracil. DFBS was sterilized via Steriflip-GP, 0.22 μm (MilliporeSigma, St. Louis, MO, USA) prior to addition to media. The erlotinib-resistant cell line HCC827ER was given as a gift from Dr. John D. Haley (Renaissance School of Medicine at Stony Brook University). HCC827ER was maintained in RPMI-1640 Medium (Thermo Fisher Scientific, Waltham, MA, USA) supplemented with 10% FBS and 1 μM erlotinib (MedChem Express, South Brunswick Township, NJ, USA).

The luciferase-expressing lung cancer line, A549-Luc, was generated by transfecting the A549 line with a luciferase-expressing lentiviral construct (gift from the Y. Ma laboratory at Renaissance School of Medicine at Stony Brook University). After transfection with the lentiviral construct, the line was maintained in Ham’s F-12K (Kaighn’s) Medium supplemented with 10% FBS and 1 ug/ml puromycin (MilliporeSigma, St. Louis, MO, USA). Luciferase expression was confirmed via IVIS before the cell line was used for *in vivo* experiments.

### Transfection

Cell lines were transfected with Oligofectamine™ (Thermo Fisher Scientific, Waltham, MA, USA) and oligonucelotides as previously described.^34,46^ In brief, A549, Calu-1, A549 (5-FU), and HCC827ER cell lines were seeded onto 6-well plates at a cell density of 1 × 10^5^ cells/well. 24 hours later, cells were transfected with Oligofectamine™ and their respective oligonucleotides according to the manufacturer’s protocol. 24 hours post-transfection, 5-FU was added to their respective wells. All oligonucleotides were transfected at a concentration of 50 nM. 5-FU was either added at 450 nM, a concentration equivalent to 5-FU in 5-FU-miR-129, or at 11.67 μM, the IC_50_ of 5-FU in A549.

Vehicle-free transfections were performed as previously described. In brief, cell lines were seeded onto 6-well plates at 1 × 10^5^ cells/well. 24 hours after seeding, the cells were transfected with their respective oligonucleotides by directly adding the oligonucleotides to their respective wells. 24 hours post-transfection, media was changed to fresh media supplemented with 10% DFBS and 5-FU was added to their respective wells.

5-FU-miR-129 was purchased from Horizon Discovery (Horizon Discovery, Cambridge, UK) as previously described. In brief, hsa-miR-129-5p’s sequence was modified by substituting uracil residues with 5-FU. Hsa-miR-129-3p’s sequence was left unmodified to avoid potential off-target effects and to preserve miRNA function. The two strands were then annealed according to the manufacturer’s protocol. A non-specific scramble miRNA, Pre-miR™ Negative Control #2 (Thermo Fisher Scientific, Waltham, MA, USA) was used to represent the negative control. siBCL2 (Smart Pool) and siHMGB1 (Smart Pool) were purchased from Horizon Discovery (Horizon Discovery, Cambridge, UK). Pre-miR-129-5p was purchased from Thermo Fisher Scientific (Thermo Fisher Scientific, Waltham, MA, USA).

### Cell Proliferation Analysis

For treatment studies with vehicle (Oligofectamine™), cell lines were seeded onto 6-well plates at 1 × 10^5^ cells/well and transfected with their respective oligonucleotides as described above. 24 hours post-transfection, cells were trypsinized and re-seeded onto 96-well plates at 1000 cells/well in triplicate per condition in fresh media supplemented with 10% DFBS. 5-FU was also added to their respective wells. Cell viability was measured using WST-1 Cell Proliferation Reagent (MilliporeSigma, St. Louis, MO, USA) according to the manufacturer’s protocol. In brief, cells were incubated in 100 μl fresh media supplemented with 10% DFBS and 10 μl WST-1 for 1 hour at 37°C. After incubation, absorbance was measured at 450 nm and 630 nm with O.D. calculated by subtracting absorbance at 630 nm from absorbance at 450 nm. Absorbances were then normalized to the negative control. Cell viability was measured at their respective time points (1, 3, and 6 days post-transfection).

For treatment studies without a transfection vehicle, cell lines were seeded onto 96-well plates at 1000 cells/well in triplicate per condition. 24 hours after seeding cells onto plates, media was changed to oligonucleotides mixed with fresh media supplemented with 10% DFBS for their respective conditions. 24 hours post-transfection, media was changed to fresh media supplemented with 10% DFBS and 5-FU was added to their respective wells. Cell viability was measured using WST-1 Cell Proliferation Reagent as described above.

### Cytotoxicity Assay

Cytotoxicity of 5-FU-miR-129 was measured by transfecting cell lines as described above with varying concentrations of 5-FU-miR-129 in 6-well plates. Cells were then re-seeded onto 96-well plates. 6 days post-transfection, cell viability was measured with WST-1 Cell Proliferation Reagent as described above. IC_50_ values were calculated using GraphPad Prism 9 (GraphPad Software, San Diego, CA, USA).

Cytotoxicities of chemotherapy drugs 5-FU, cisplatin (MilliporeSigma, St. Louis, MO, USA) and erlotnib were measured by seeding cells onto 96-well plates at 1000 cells/well per drug concentration. 24 hours post-seeding, media was changed to fresh media supplemented with 10% DFBS and their respective concentrations of drugs. 48 hours post-treatment, media was changed with fresh media supplemented with 10% DFBS. 6 days post-treatment, cell viability was measured with WST-1 Cell Proliferation Reagent as described above. IC_50_ values were calculated using GraphPad Prism 9.

### Western Immunoblot Analysis

Cell lines were seeded and transfected with or without a transfection vehicle (Oligofectamine™) in 6-well plates with their respective conditions as described above. 3 days post-transfection, cells were lysed with a mixture of RIPA buffer (MilliporeSigma, St. Louis, MO, USA) and protease inhibitor cocktail (MilliporeSigma, St. Louis, MO, USA) and protein samples were collected for western immunoblot analysis. Proteins were probed with mouse anti-Bcl-2 antibody (Thermo Fisher Scientific, MA511757, 1:100), rabbit anti-HMGB1 antibody (Cell Signaling, 3935S, 1:500), mouse anti-p21 antibody (Cell Signaling, 2946S, 1:500), rabbit anti-p53 antibody (Thermo Fisher Scientific, BS-8687R, 1:500), rabbit anti-LC3B (Cell Signaling, 3868S, 1:1000), or mouse anti-β-actin antibody (MilliporeSigma, A5441, 1:10,000,000). Bcl-2, p21, p53 and β-actin primary antibodies were diluted in 5% milk (Bio-Rad, Hercules, CA, USA) in TBST. HMGB1, and LC3B primary antibodies were diluted in 5% BSA (MilliporeSigma, St. Louis, MO, USA) in TBST. After staining with primary antibodies, proteins were probed with secondary antibodies goat anti-mouse-HRP (Bio-Rad, 1706516, 1:5000) or goat anti-rabbit-HRP (Bio-Rad, 1721019, 1:5000) with their respective primary antibodies. Proteins were then treated with SuperSignal™ West Pico PLUS Chemiluminescent Substrate (Thermo Fisher Scientific, Waltham, MA, USA) and visualized with a LI-COR Biosciences Odyssey Fc imaging system. Protein bands were quantified with Image Studio, Version 5.2.5 (LI-COR Biosciences, Lincoln, NE, USA).

### Autophagy Analysis

3 days post-transfection, cells were washed with PBS and media was changed to either fresh media or Earle’s Balanced Salt Solution (EBSS) (Thermo Fisher Scientific, Waltham, MA, USA) in order to either inhibit (Negative Control (+Serum)) or induce autophagy (Negative Control (Serum-Starved), respectively. Cells were then incubated for 4 hours at 37°C before protein samples were collected for western immunoblot analysis as described above. Autophagy was measured via the expression ratio of LC3B-II/LC3B-I.

### Cell Cycle Analysis

3 days post-transfection, cells were resuspended at 0.5 to 1 × 10^6^ cells/ml in Krishan modified buffer supplemented with 0.02 mg/ml RNase H (Thermo Fisher Scientific, Waltham, MA, USA) and 0.05 mg/ml propidium iodide (MilliporeSigma, St. Louis, MO, USA). Cells were then analyzed by flow cytometry via CytoFLEX Flow Cytometer (Beckman Coulter, Brea, CA, USA) and results were analyzed by Modfit LT Software (BD Biosciences, Sparks, MD, USA).

### Apoptosis Assay

3 days post-transfection, cells were resuspended in annexin-binding buffer (Thermo Fisher Scientific, Waltham, MA, USA). Cells were then stained with Annexin V (Thermo Fisher Scientific, Waltham, MA, USA) and propidium iodide (MilliporeSigma, St. Louis, MO, USA). Stained cells were analyzed by flow cytometry via CytoFLEX Flow Cytometer (Beckman Coulter, Brea, CA, USA).

### Cell Death Profiler Assay

3 days post-transfection, total RNA was extracted from transfected cells using TRIzol reagent (Thermo Fisher Scientific, Waltham, MA, USA). From the extracted total RNA, cDNA was prepared using QIAGEN’s RT^2^ First Strand Kit (QIAGEN, Hilden, Germany). RT-PCR was performed using QIAGEN’s RT^2^ Profiler™ PCR Array Human Cell Death PathwayFinder (QIAGEN, Hilden, Germany). RT-PCR data was analyzed and visualized for pathway analysis by GeneSpring GX 14.9 (Agilent Technologies, Santa Clara, CA, USA) and by Enrichr (Icahn School of Medicine, Mount Sinai, NY, USA).^43–45^

### Mouse Non-Small Cell Lung Cancer Metastasis Model

All animal procedures were approved by the Stony Brook University Institutional Animal Care and Use Committee (IACUC). Non-obese diabetic (NOD)/severe combined immunodeficiency (SCID) mice were purchased from The Jackson Laboratory (The Jackson Laboratory, Bar Harbor, ME, USA). 8-week-old NOD/SCID mice (2 – 3 males and 2 – 3 females per treatment group) were injected with 1 × 10^7^ A549 (+Luc) cells via intravenous (IV) tail vein injection. Mice were then divided into four groups (control, 5-FU, gemcitabine, and 5-FU-miR-129). Control and 5-FU-miR-129 groups were treated with either 75 μg of PEI-MAX (control) (Polysciences, Warrington, PA, USA) or 75 μg PEI-MAX + 75 μg 5-FU-miR-129 (5-FU-miR-129). Average mouse weight ≈ 20 g, 75 μg injection/20 g weight = 3.75 mg/kg. 5-FU and gemcitabine (Gem) groups were treated with either 50 mg/kg 5-FU (5-FU) or 50 mg/kg Gem (MilliporeSigma, St. Louis, MO, USA). In the first treatment session (Treatment 1), control and 5-FU-miR-129 groups were treated via IV tail vein injection every other day for 2 weeks (eight times), and 5-FU and gemcitabine groups were treated via intraperitoneal (IP) injection every 3 days for 2 weeks (four times). In the second treatment session (Treatment 2), control and 5-FU-miR-129 groups were treated via IP injection every other day for 1 week (four times), and 5-FU and Gem groups were treated via IP injection every 3 days for 1 week (two times). Treatment sessions started at 2 days post-injection of A549 (+Luc) cells (Day 2, Treatment 1) and 33 days post-injection (Day 33, Treatment 2). 10 minutes prior to measuring luciferase expression via luciferase activity, mice were injected with RediJect D-Luciferin (PerkinElmer, Waltham, MA, USA). Luciferase expression was used to measure tumor growth using the IVIS Spectrum *In Vivo* Imaging System (IVIS) (PerkinElmer, Waltham, MA, USA). Tumor growth was calculated as a function of (total flux at time of measurement [p/s])/(total flux at initial measurement [p/s]). Mice were monitored daily and euthanized according to IACUC guidelines to determine survival time.

### Statistical Analysis

All experiments were performed at least three times, and all statistical analyses was performed using GraphPad Prism 9 (GraphPad Software, San Diego, CA, USA). Statistical significance between two groups was determined using Student’s t-test. Data is presented as mean ± SD. Statistical significance was presented either as described in the figure legends or with the following asterisks (*p < 0.05; **p < 0.01; *** p < 0.001; **** p < 0.0001).

## Acknowledgements

We would like to thank the Dr. Yupo Ma lab at the Renaissance School of Medicine at Stony Brook University for providing the luciferase-expressing lentivirus. This research was supported by, in part, the USA National Institute of Health/National Cancer Institute (NIH/NCI), grant number R01CA1550197098 (J. Ju), Curamir Therapeutics Inc. (J. Ju), and the VA Merit Award, grant number BX005260-01 (J. Ju).

## Declaration of Interests

A.F. and J.J. have filed a patent for 5-FU modified miRNA mimetics. J.J. is a scientific co-founder of Curamir Therapeutics. The remaining authors declare no competing interests.

## Author Contributions

Conceptualization, G.H., and J.J; Methodology, G.H. and J.J; Investigation, G.H., J.G.Y., A.F., H.F.; Writing – Original Draft, G.H. and J.J.; Writing – Reviewing & Editing, G.H., J.G.Y., A.F., J.D.H., and J.J.; Supervision, J.J.; Project Administration, J.J.; Funding Acquisition, J.J.

## Notes

### Summary of Updates

There was an error during figure assembly with Supplemental Figure S3. Supplemental Figure S3 has been updated to fix this error.

