## Supplemental Figures and Tables for "Development of a 5-FU Modified miR-129 Mimic as a Therapeutic for Non-Small Cell Lung Cancer"

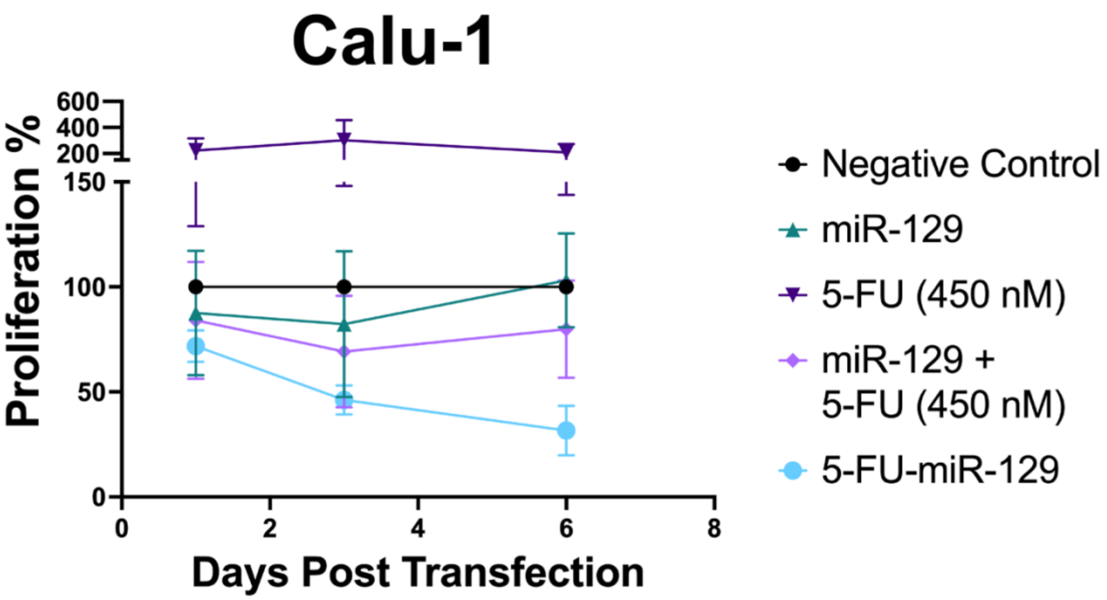


**Figure S1: WST-1 Assay 1, 3, and 6-Days Post-Transfection in Calu-1 cells.** Proliferation was measured by WST-1 assay 1, 3, and 6-days post-transfection in Calu-1 cells. By 3 days post-transfection, miR-129 and 5-FU-miR-129 were observed to inhibit proliferation in Calu-1. Data are represented as mean ± SD.

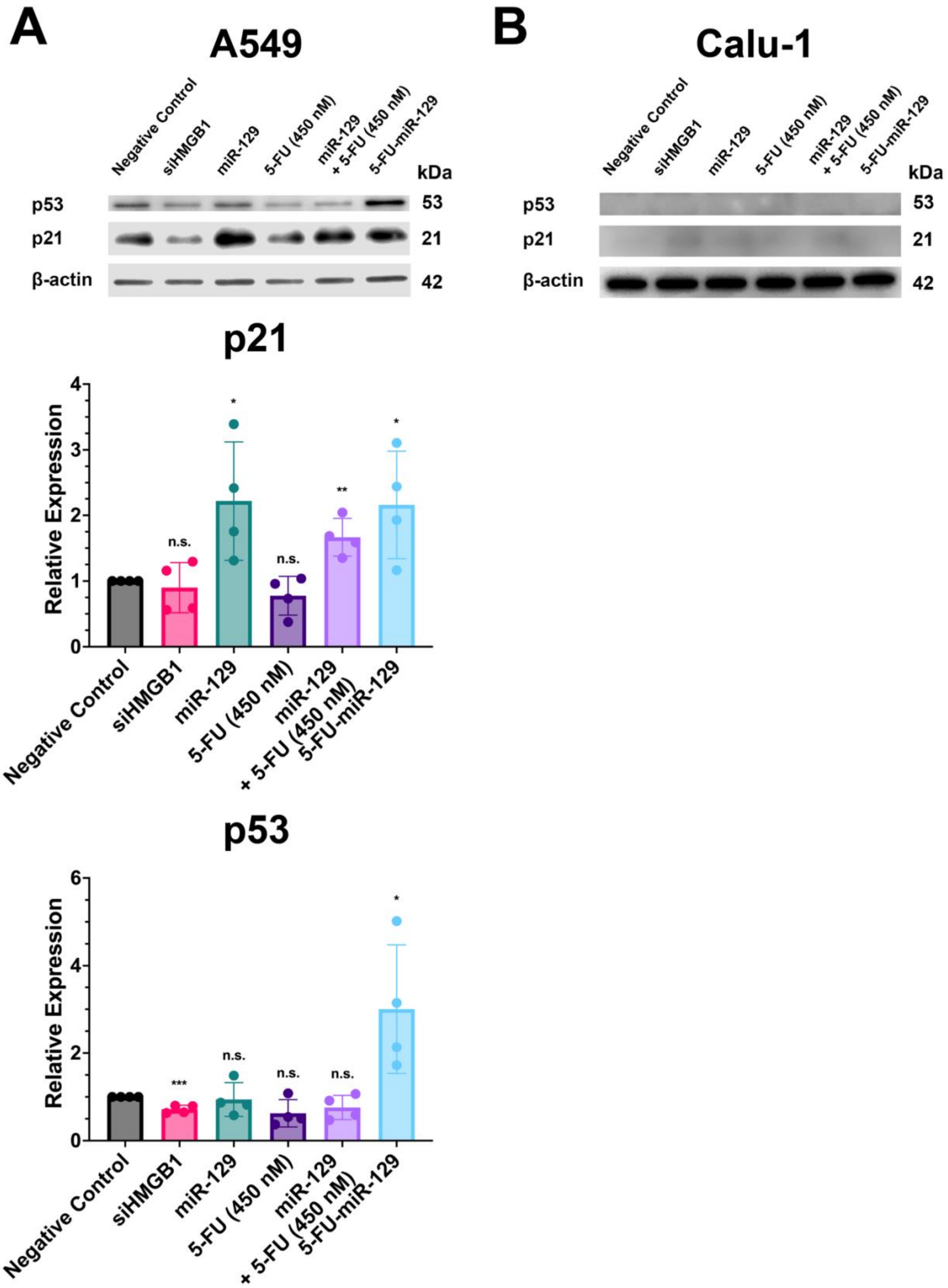
A
A549
Negative Control
siHMGB1
miR-129
5-FU (450 nM)
miR-129
+ 5-FU (450 nM)
5-FU-miR-129
kDa
p53
p21
β-actin
53
21
42
B
Calu-1
p21
Relative Expression
n.s.
*
**
p53
***
0
1
2
3
4
6

**Figure S2: 5-FU-miR-129 Promotes Expression of p21 and p53 in A549, But Not in Calu-1 cells.** (A) Transfecting cells with 5-FU-miR-129 was found to increase the expression of p21 and p53 in A549 cells, as seen by an increase in p21 and p53 expression ($p = 0.0301$ and $p = 0.0343$, respectively). An increase in p21 expression was also observed in cells transfected with either miR-129 ($p = 0.0358$) or miR-129 + 5-FU ($p = 0.0035$). *$p < 0.05$, **$p < 0.01$ (n = 4). A decrease in p53 expression was also observed in cells treated with siHMGB1 ($p = 0.0007$). *$p < 0.05$, ***$p < 0.001$ (n = 4). (B) Significant changes in the expression of p21 and p53 were not observed in Calu-1. All transfections were performed with 50 nM of negative control, siRNA controls, miR-129, or 5-FU-miR-129. Cells were also treated with 450 nM of 5-FU to mimic the equivalent concentration of 5-FU in 5-FU-miR-129. p21 and p53 western blots in A549 and Calu-1 are from the same blots shown, respectively, in Figure 2. Data are represented as mean ± SD.

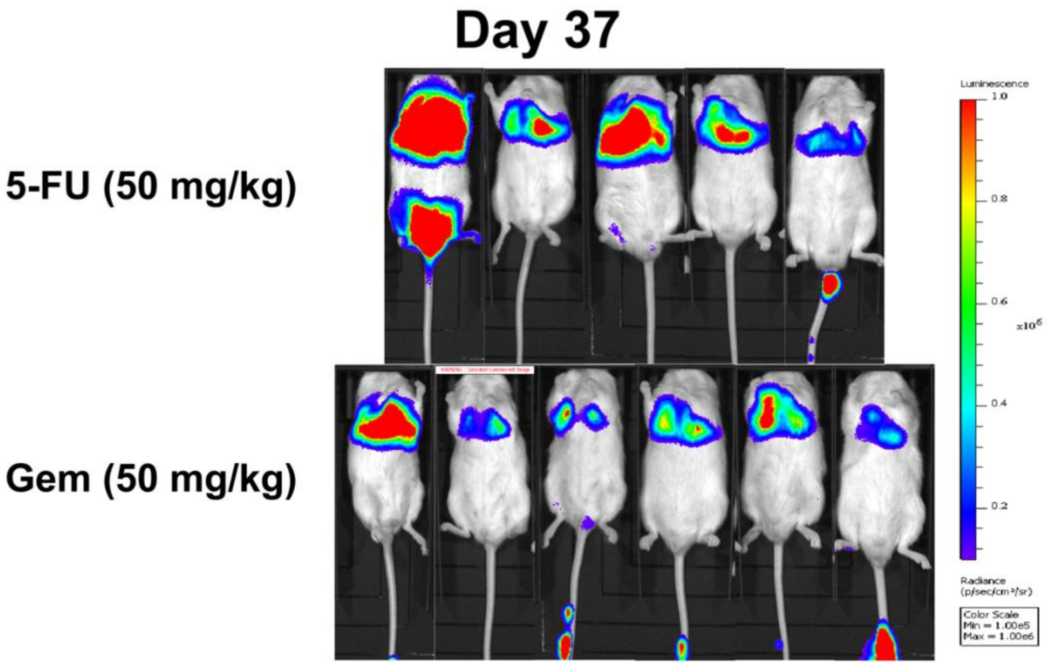


**Figure S3: Luciferase Expression of 5-FU and Gem Treated Groups 37 Days Post NSCLC Injection.** The IVIS imaging system was also used to visualize NSCLC tumor growth in 5-FU and Gem treated groups 37 days post-injection with A549 (Luc) cells (1 x $10^7$ cells per mouse).

| Treatment | $IC_{50}$ (nM) |
| --- | --- |
| 5-FU | 11,665 |
| Gemcitabine | 498.48[a] |
| Cisplatin | 11,370 |
| 5-FU-miR-129 | 13.33 |

**Table S1: $IC_{50}$ Values of Chemotherapies and 5-FU-miR-129 in A549 Cells.** [a]The $IC_{50}$ of gemcitabine was determined by previously reported values.[42]

| Upregulated in A549 | | | Downregulated in A549 | | |
|---|---|---|---|---|---|
| Gene | Fold Change | Pathway | Gene | Fold Change | Pathway |
| APAF | 2.28 | apoptosis | BCL2 | 0.25 | apoptosis, autophagy |
| APP | 2.06 | autophagy | BMF | 0.34 | necrosis |
| BCL2A1 | 2.26 | apoptosis | C1orf159 | 0.5 | necrosis |
| BIRC3 | 4.15 | apoptosis | CD40LG | 0.41 | apoptosis |
| CASP1 | 2.21 | apoptosis | DEFB1 | 0.33 | necrosis |
| CASP7 | 2.28 | apoptosis | ESR1 | 0.12 | autophagy |
| CD40 | 2.43 | apoptosis | PARP2 | 0.49 | necrosis |
| CFLAR | 2.3 | apoptosis | TNFRSF11B | 0.47 | apoptosis |
| CTSB | 2 | autophagy | ULK1 | 0.47 | autophagy |
| CTSS | 2.5 | autophagy | | | |
| CYLD | 2.3 | apoptosis, necrosis | | | |
| DPYSL4 | 14.02 | necrosis | | | |
| EIF5B | 2.24 | necrosis | | | |
| FAS | 15.33 | apoptosis, autophagy | | | |
| GADD45A | 2.32 | apoptosis | | | |
| GALNT5 | 6.11 | necrosis | | | |
| SYCP2 | 2.85 | apoptosis, necrosis | | | |

**Table S2: Observed Changes in mRNA Expression in Genes Related to Cell Death by 5-FU-miR-129 in A549 Cells Identified by RT$^2$ Profiler Array.**

| Upregulated in Calu-1 | | | Downregulated in Calu-1 | | |
|---|---|---|---|---|---|
| Gene | Fold Change | Pathway | Gene | Fold Change | Pathway |
| ATG3 | 2.02 | autophagy | APAF1 | 0.31 | autophagy |
| BCL2A1 | 7.15 | apoptosis | CCDC103 | 0.5 | necrosis |
| BIRC2 | 2.26 | apoptosis | GALNT5 | 0.48 | necrosis |
| CASP9 | 2.14 | apoptosis | SPATA2 | 0.41 | apoptosis, necrosis |
| CYLD | 2.09 | apoptosis | TNFRSF1A | 0.42 | apoptosis, necrosis |
| DPYSL4 | 2.13 | necrosis | ULK1 | 0.46 | autophagy |
| FAS | 2.07 | apoptosis, autophagy | | | |
| FOXI1 | 3.43 | necrosis | | | |
| GADD45A | 2.16 | apoptosis | | | |
| HSPBAP1 | 2.03 | necrosis | | | |
| IRGM | 4.52 | autophagy | | | |
| KCNIP1 | 6.41 | necrosis | | | |
| RAB25 | 4.37 | necrosis | | | |

**Table S3: Observed Changes in mRNA Expression in Genes Related to Cell Death by 5-FU-miR-129 in Calu-1 Cells Identified by $RT^2$ Profiler Array.**
